# Scalp microbiome differences in subjects with self-reported hair loss: A quantitative approach to microbial dysbiosis

**DOI:** 10.1101/2025.03.11.642466

**Authors:** Judy Malas, Sebastian Reczek, Imani Porter, Yolanda M. Lenzy, Victoria Barbosa, Crystal Porter, Jarrad Hampton-Marcell

## Abstract

**Objective:** Hair loss is a common issue that affects a large proportion of the population, leading to lower self-confidence and quality of life. Microbial dysbiosis of the scalp has been shown to be associated with several different disorders leading to hair loss. Though several “microbiome friendly” cosmetic treatments are currently on the market, there is no agreement on the best technique for assessing dysbiosis leading to a lack of scientific rigor for quantifying the effective treatments. To help address this, the association between self-perceived hair loss and the scalp microbiome in an African-American cohort (n=36) was investigated.

**Methods:** Using a self-controlled design, swabs were collected from both “sparse” and “normal” scalp sites. The scalp microbiome was characterized via 16S rRNA gene sequencing and a dysbiosis score was calculated based on the proportion of all taxa within the samples. Further, we identified the taxa that contributed most to abnormal or dysbiotic hair sites using a machine learning random forest classifier and a negative binomial mixed effects model.

**Results:** The dysbiosis index is sensitive to participants self-assessment of hair loss and interindividual variation. We found a core set of operational taxonomic units (OTUs) assigned to 7 genera that significantly contributed to increased scalp dysbiosis.

**Conclusion:** This study demonstrates that self-perceived hair loss is associated with significant and measurable alterations in the scalp microbiome using, making the reported dysbiosis index a practical tool that may be used to assess microbiome changes following cosmetic or medical interventions for hair loss and other microbiome-associated disorders.

## Introduction

Hair loss is a pervasive concern with a multifactorial etiology, encompassing factors such as stress, nutritional deficiencies, harsh styling practices, and underlying medical conditions[1]. Androgenetic alopecia, the most common form, affects a significant portion of the global population, with prevalence varying by ethnic group [2–4]. Given the central role of hair in aesthetic standards and self-perception, hair loss is frequently associated with diminished self-confidence and a reduced quality of life [5], driving individuals to seek both medical and cosmetic interventions [6].

The microbiome has emerged as a critical factor influencing skin and scalp health [5]. The scalp presents a unique ecological niche, characterized by high density of hair follicles and sebaceous glands, which support a distinct microbial community. Furthermore, individual hair follicles harbor their own microbiota, with different bacterial species exhibiting varying capacities to adhere to and colonize hair shafts [7, 8].

This has led to growing interest in the association between microbial dysbiosis and alopecia. The term “dysbiosis” generally describes a departure from a healthy microbial state, though its application ranges from broad shifts in community structure to alterations in specific taxa[9, 10]. Some studies have proposed specific ratios of abundant genera like *Cutibacterium, Staphylococcus*, and *Corynebacterium* as markers of scalp dysbiosis [11–14], while others argue that a holistic view of community composition is more accurate [15, 16]. While the causal relationship between these microbial shifts and hair loss remains to be fully elucidated, understanding these associations is crucial for developing targeted symptom management and therapeutic strategies.

The cosmetic industry has taken note, with an increasing number of products marketed as “microbiome-friendly.” Cosmetics and personal care products are known to exert significant and lasting effects on microbial composition [17, 18]. However, a major challenge in developing effective products lies in the objective assessment of dysbiosis and product efficacy. A standardized, quantitative metric to evaluate scalp dysbiosis is therefore urgently needed to guide product development and claims substantiation. Critically, in both clinical and cosmetic contexts, a persistent challenge has been the translation of objective scientific measurements into tangible consumer benefits. It is exceptionally rare for a microbiological metric to be directly correlated with a subjective consumer perception, such as the observation of hair thinning or loss.

While prior investigations have primarily focused on specific alopecia subtypes, the present study examines hair loss more broadly within an African American cohort reporting general hair loss. Using a self-controlled design, we collected scalp swabs from adjacent “sparse” and “normal” sites (no perceived hair loss) from 36 individuals and characterized the microbiota via 16S rRNA gene sequencing. We hypothesized that sites of self-assessed hair loss would exhibit distinct microbiome compositions compared to unaffected sites on the same scalp. Our results not only confirm this hypothesis but, more significantly, demonstrate that our newly developed dysbiosis index shows a promising correlation with perceived hair loss. This suggests the potential of this index as a valuable tool to quantitatively bridge the gap between microbial ecology and consumer experience in the assessment of hair loss.

## Methods

### Recruiting and Sample Collection

Adults with self-reported hair loss were recruited for the study without restrictions in ethnic background. Exclusion criteria included participants with complete hair loss, the presence of open sores, drainage and or injury to the scalp, and cancer or chemotherapy treatment. Participants were instructed to use a sterile swab (Puritan catalog # BD 25-806 10WC) to collect two samples: one from an afflicted (hair-loss) site and one from a normal site, swabbing the areas for 10 seconds each to collect microbial biomass. All qualifying participants completed a hair density scale; a questionnaire where participants rated their degree of hair loss using an anchored line scale from 1 to 10. An index score of 1 represented a participant whose scalp was readily viewable, and 10 represented a scalp completely covered by hair fibers [19, 20]. Participants also completed a stress scale, which contained anchored line scales where they indicated their level of stress on a scale from 1 – 10 [21]. Study protocols were approved and overseen by The University of Chicago Biological Sciences Division institutional review board, Chicago, Illinois (protocol number IRB16-1541). A signed consent was obtained from all participants prior to sampling and questionnaire completion.

### Library preparation and sequencing

Swabbed samples were placed inside of a sterile Whirl Pak® bag and mailed to the testing facility. Upon arrival, samples were stored in a -80°C freezer until DNA extraction. Genomic DNA was extracted using the recommended protocols of the Earth Microbiome Project (EMP) and American Gut Project [22, 23]. The V4 region of the 16S rRNA gene (515F-806R) was amplified. Barcoded primer sets were adapted for the Illumina MiSeq System as described by [24]. The reverse amplification primer also contained a twelve-base barcode sequence that supported pooling of up to 2,167 different samples in each lane. Each polymerase chain reaction (PCR) for gene amplification contained 12ul of MoBio PCR Water (Certified DNA-free), 10ul of 5 Prime HotMasterMix (1x), 1ul of Forward Primer (5uM concentration, 200pM final), 1ul Golay Barcode Tagged Reverse Primer (5uM concentration, 200pM final), and 1ul of genomic DNA. The PCR conditions are as follows: 94 °C for 3 minutes to denature the DNA, with 35 cycles at 94 °C for 45 seconds, 50 °C for 60 seconds and 72 °C for 90 seconds, with a final extension of 10 minutes at 72 °C. Following PCR, amplicons were quantified using PicoGreen (Invitrogen) and a plate reader. Once quantified, different volumes of each of the products were pooled into a single tube so that each amplicon was represented equally. This pool was then cleaned using the UltraClean® PCR Clean-Up Kit (MoBio) and quantified using Qubit (Invitrogen). After quantification, the molarity of the pool was determined and diluted to 2nM, denatured, and then diluted to a final concentration of 2pM with a 30% PhiX spike for sequencing on the Illumina MiSeq System.

### Alpha and beta diversity

All statistical analyses were performed using R v4.4.2. Sequences were quality filtered and then assigned to corresponding samples based on their 12bp error-correcting Golay barcodes using QIIME v1.9.1 [25]. Operational taxonomic units (OTUs) were defined and assigned taxonomy by clustering sequences against 97% OTU reference data set from the Greengenes database [26, 27]. Alpha diversity measures were generated using ‘microbiome’ package v1.28.0 [28]. Pearson correlations were calculated to determine the linear relationship between age and alpha diversity. Age groups were also analyzed categorically by splitting the samples into roughly equal “old” and “young” group based the median participant age; a Wilcoxon rank-sum test was conducted to evaluate differences in alpha diversity between the two age groups. Beta diversity was calculated on Aitchison distances (Euclidean distances of center log ratio count data) to compare microbial composition between age groups and afflicted or normal hair sites. OTU count data were transformed using center log ratio transformation in the R package ‘microbiome’. Following data transformation, a principal components analysis (PCA) was conducted with R package ‘phyloseq’ [29]. A Permutational multivariate analysis of variance (PERMANOVA) test was conducted within the R package ‘Vegan’ (v2.6-4). PERMANOVA tests and p values were calculated based on 10,000 permutations.

### Dysbiosis Index

Dysbiosis scores were calculated using the R package ‘dysbiosisR’ v1.0.4 (https://github.com/microsud/dysbiosisR). The dysbiosis index was defined using a modified version of the method reported in [30]. Here, the Aitchison dissimilarity distance matrix for the overall community was used rather than Euclidean distance based on quantitative PCR of select taxa as in AhShawaqfeh et al. (2017). Each microbial dysbiosis score is calculated as the difference between the Aitchison distance between the test sample and the healthy class centroid and the Atichison distance between the test sample and the diseased class centroid. In other words, the dysbiosis score for each sample measures its closeness to the group (hair-loss afflicted or normal) mean. The dysbiosis score D of the test sample z, is defined as

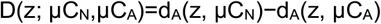

where μC_N_ and μC_A_ are the centroids (mean composition) of the normal and afflicted samples, respectively and d_A_ is the Aitchison distance either between the test sample and the μC_N_ or μC_A_. A dysbiosis score of 0 indicates that the test sample is at equal distance from the center of both class (afflicted and normal) centroid, and higher scores indicate more deviation from normal. Calculated dysbiosis scores were classified into two groups of either dysbiosis (score > 0) or non-dysbiotic (score < 0). Code used to calculate dysbiosis scores along with all microbial analyses conducted is publicly available (https://github.com/jud-m/Defining-microbial-dysbiosis).To assess the performance of the dysbiosis score as a tool for distinguishing normal and afflicted (hair loss) scalp sites, a receiver operating characteristic (ROC) curve was generated using the package ‘pROC’ and the area under the ROC curve (AUC) was calculated.

### Determination of taxa contributing to dysbiosis

To determine the bacterial taxa that differentiated dysbiotic hair sites from the normal hair sites, a random forest (RF) machine learning classifier was used to determine which data features (OTUs) were most important for classifying dysbiotic samples using the R package ‘randomForest’ v4.7-1.2. The 100 most important OTUs (out of 6,009 total OTUs) for classification were considered further. In addition, because age can be a confounding variable in determining the microbial taxa associated with dysbiosis, a negative binomial mixed-effect model (NBMM) was used to model the microbiome count data using the R package ‘NBZIMM’ v1.0 [31]. The NBMM identified differentially abundant taxa in the afflicted hair sites relative to normal sites while controlling for the effect of age. As a final check, a Wilcoxon rank-sum test was conducted on the log10 abundance of the OTUs that were flagged in both the NBMM and the random forest classifier. OTUs that were not differentially abundant according to the Wilcoxon rank-sum test were excluded.

## Results

### Cohort characteristics

Most recruited participants were African American (36 of 38) and the remaining two participants were Caucasian. Caucasian participants were excluded from further analyses as there were not enough Caucasian participants to have robust data for comparison. The average age of the participants was 51.6 years (sd +/-13.5) and the median age was 52 years. The cohort included 29 female and 7 male participants. The state of the hair among participants included natural (n = 20), relaxed (n = 8), color (n = 5), keratin treatment (n = 1), and other (n = 2). None of the participants wore hair prostheses. Both scalp density (Pearson, p = 0.0019, R^2^ = -0.51) and scalp stress (Pearson, p = 0.073, R^2^ = -0.31) decreased with age but only the correlation with scalp density was statistically significant. The frequency of scalp cleansing reported by participants included monthly (n = 7), biweekly (n = 16), weekly (n = 12), and daily (n = 1). Self-reported measures of hair density and stress decreased slightly with age, but associations were not statistically significant.

### Microbial diversity

Following quality filtering, 1,720,130 microbial sequences comprising 6,009 OTUs were observed for 67 samples (35 afflicted, 32 normal). Firmicutes (50.6%), Actinobacteria (33%) and Proteobacteria (11.6%) were the most abundant phyla. *Staphylococcus, Corynebacterium*, and *Propionibacterium* were the most abundant genera (Fig. 1). Shannon alpha diversity, a measure of richness and evenness of species within a sample, was significantly higher in the afflicted hair sites (p = 0.004). Shannon alpha diversity was positively correlated with age in both afflicted and normal hair sites (R^2^ = 0.62 and 0.4, respectively; Fig. 2A). Although diversity increased with age, it increased to a greater extent in the afflicted hair sites, as reflected by the higher R^2^ value for the afflicted hair sites. When categorized into two groups, participants 52 years (median age) or older have significantly higher Shannon diversity values in the afflicted sites (Fig. 2B). Hair state (including treatments), cleansing frequency, participant sex, and self-reported stress and hair density were not significantly associated with alpha diversity.

**Figure 1.**
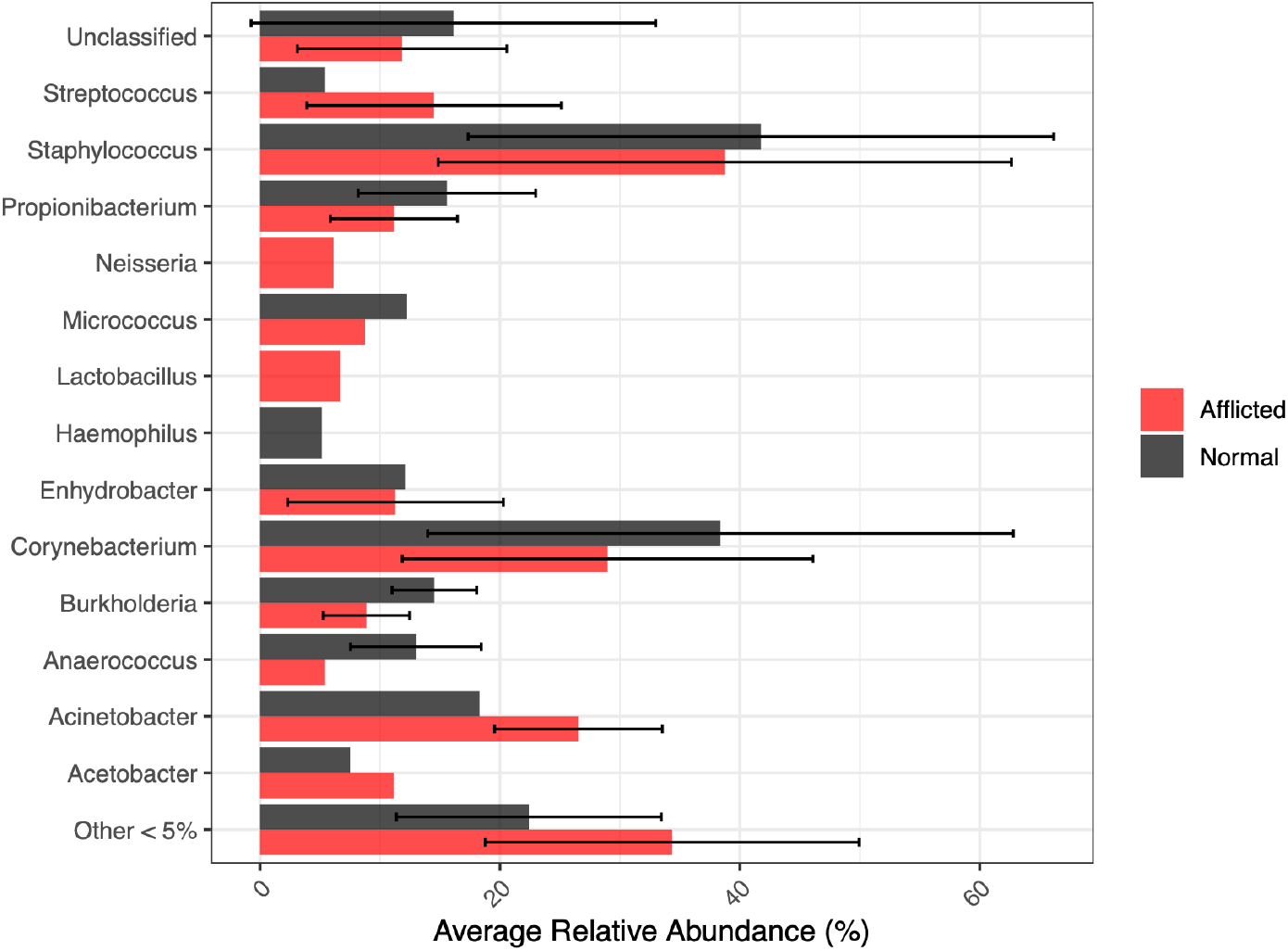
Average relative abundance of abundant genera, considering all OTUs, in afflicted and normal hair sites. Error bars show standard deviation between samples within the afflicted and normal groups.

**Figure 2.**
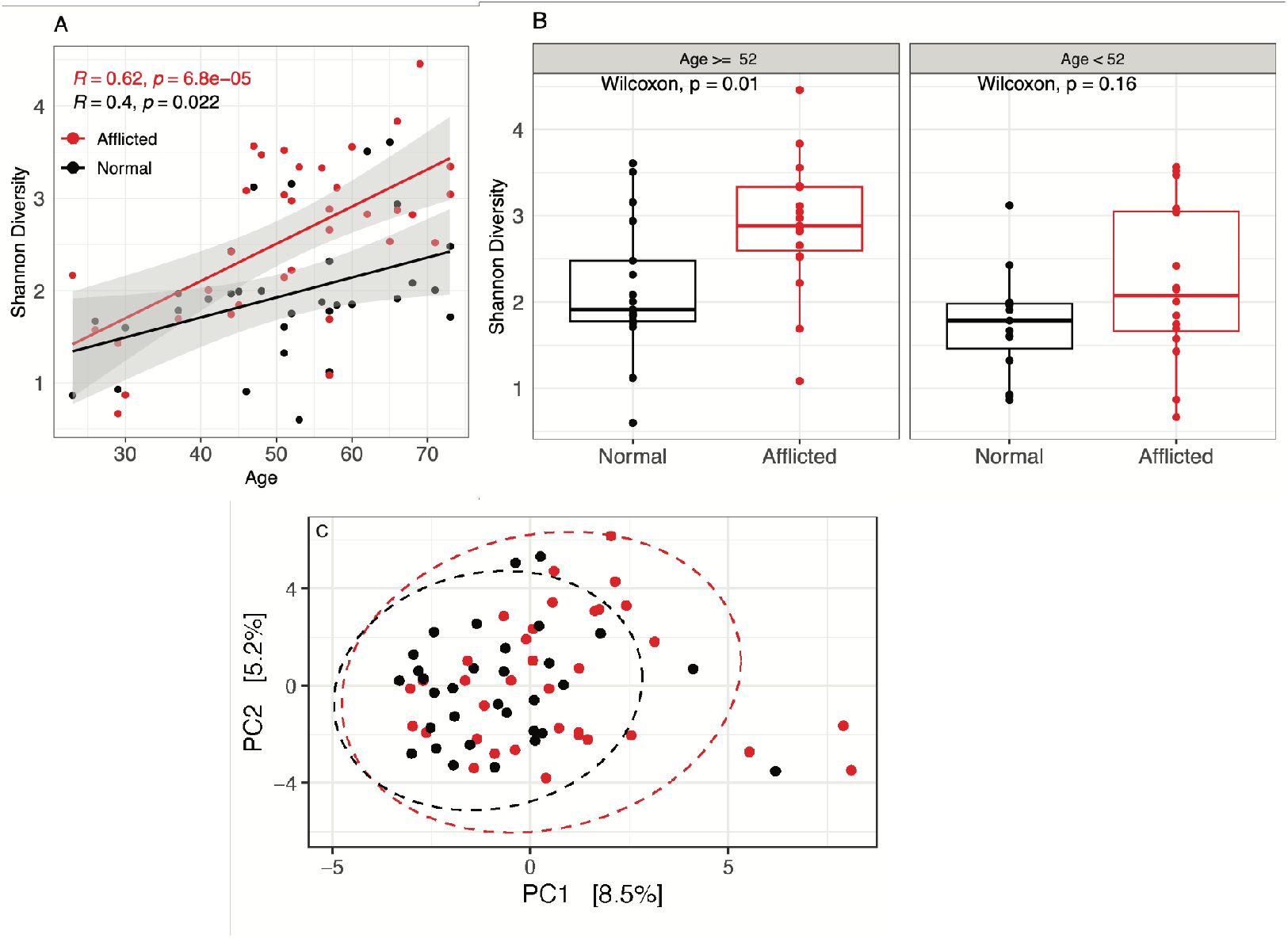
A) Linear regression of age of participants with Shannon Alpha diversity of the scalp microbiome. Red points are the afflicted sites, black points are the normal sites. B) Shannon alpha diversity of the afflicted and normal hair site. Panels split into samples from particpants greater or equal to the median age (52), or less than the median age. C) Euclidean distance PCA of center-log ratio transformed counts (Aitchson distances) of all samples.

There is significant overlap in the overall community composition between afflicted and normal hair sites (Fig. 2C). PC1 accounted for 8.5% of the variance while PC2 accounted for 5.2% of the variance in the PCA plot. This overlap can potentially be explained as a byproduct of study participants contributing both afflicted and normal samples. Despite the overlap, PERMANOVA results indicated that there is a statistically significant difference in the overall microbial community composition between afflicted and normal hair sites (p = 0.048, R^2^ = 0.0189). Notably, the result was significant even when limiting permutations within participants to account for the matched-pair design of the study using the “strata” argument of the PERMANOVA test (p= 9.9 x 10^-5^). Though participant sex was not significantly associated with alpha diversity, beta diversity was significantly different between male and female participants (p = 3×10^-4^). A PERMANOVA test was also conducted on the age groups (< 40, 40-59, 60+) and the overall microbial community was found to be significantly different between groups (p = 0.0011), suggesting both age and sex play an important role in the overall microbial community composition.

### Dysbiosis Index

A modified dysbiosis index was used to calculate a dysbiosis score for each sample. Effectively, the index determines whether the microbial composition was of each individual sample closer in composition to the afflicted samples versus or the normal samples. Scores < 0 (closer to normal) were considered non-dysbiotic and scores > 0 (closer to afflicted) were considered dysbiotic. It was determined that the majority, but not all, sites of hair loss fall into the dysbiotic classification (Fig. 3). Based on the threshold set, 24 of 35 afflicted samples were classified as dysbiotic (68.5%) and 30 of 32 samples normal samples as non-dysbiotic (93.75%).

**Figure 3.**
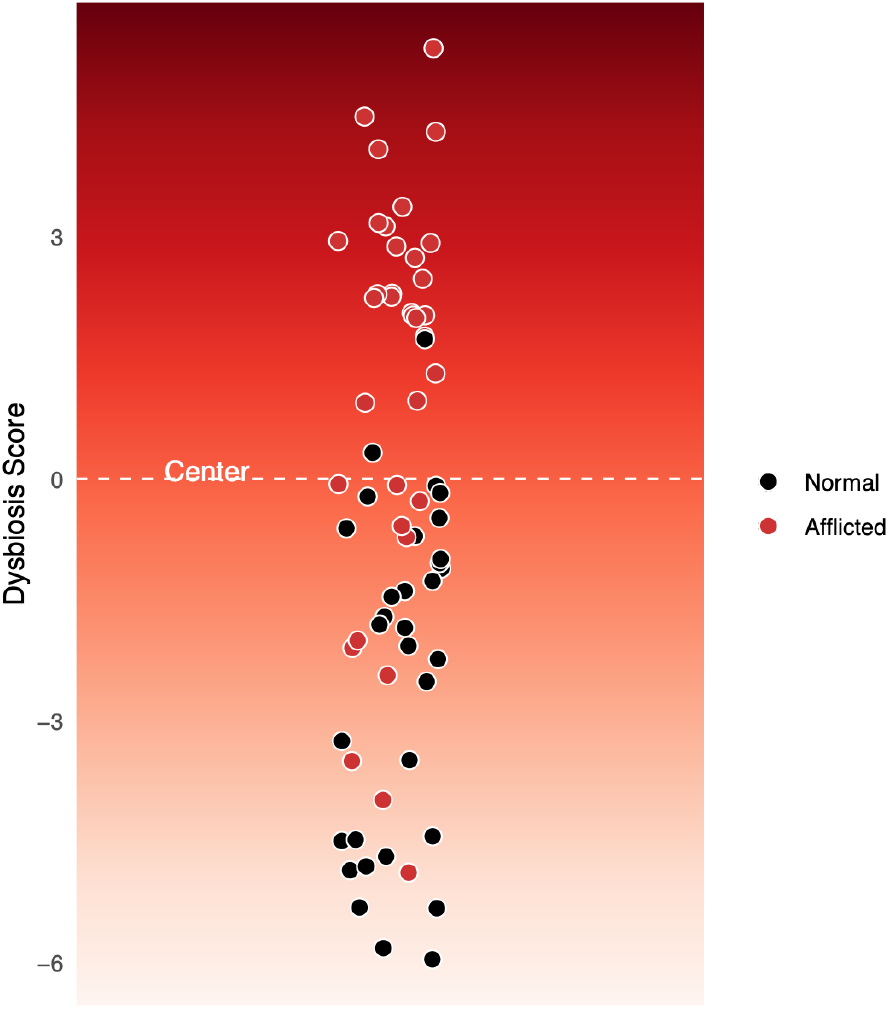
Dysbiosis scores plotted with threshold (white dotted line). A dysbiosis score of 0 indicates that the test sample is at equal distance from the center of both class (afflicted and normal) centroid, and higher scores indicate more deviation from normal.

### Core taxa contributing to dysbiosis

A RF machine learning classifier was used to determine the most important OTUs for distinguishing the dysbiotic samples. Assuming all sites of hair loss are dysbiotic, the overall error rate for the RF was 20.9%. The model was better at classifying normal sites; dysbiotic sites were classified to an accuracy of 57.6%, while the normal sites were correctly classified in 92.6% of samples. Twenty-seven unique genera were assigned to the 100 most important OTUs for classification (Fig. S1). The mean decrease accuracy (MDA), a measure that indicates the percentage decrease in the random forest model accuracy when a specific OTU is excluded, was >10% for OTUs assigned to the genera *Streptococcus* (66%), *Corynebacterium* (34.2%), *Anaerococcus* (15.8%), *Haemophilus* (14.3%), *Peptoniphilus* (12.5%), *Staphylococcus* (11.9%), *Actinomyces* (11.4%), and *Rothia* (10.5%).

To control for the potential confounding effect of age, a NBMM was used to model the microbiome count data using including age a random effect variable and scalp condition (afflicted or normal) as the fixed effect. This approach removes microbial variation associated with age prior to analyzing microbial taxa differentially abundant between sites. A NBMM was generated and independently tested for each of the 6009 OTUs. A total of 100 OTUs belonging to 29 unique genera were found to be differentially abundant in the afflicted sites compared to the normal sites, the majority of which (83%) were more abundant in the afflicted sites (Fig S2). The top genera *Corynebacterium* (18 OTUs), *Streptococcus* (17 OTUs), *Acinetobacter* (8 OTUs), *Anaerococcus* (8 OTUs), *Staphylococcus* (7 OTUs), and *Enhydrobacter* (3 OTUs) were assigned to more than 2 OTUs (Fig S2). Though species level assignments were not possible for the majority of the OTUs, several species were assigned including: *Staphylococcus epidermidis, Propionibacterium acnes, Corynebacterium durum, Lactobacillus inners, Paracoccus aminovorans, Veillonella dispar. S. epidermidis* and *P. acnes* decreased in the afflicted sites, while the remaining species increased in the afflicted sites. Two different OTUs were assigned to the species *Rothia dentocariosa*; one OTU of the species increased, and one decreased in the afflicted sites.

There were 42 OTUs in common from the RF and NBMM tests. Of these 42, OTUs, only 20 assigned to 7 genera were found to be differentially abundant after conducting a Wilcoxon-rank-sum test (Fig 4). The majority of these OTUs were more abundant in the afflicted site, in line with the increased diversity observed in the afflicted sites (Fig 2A-B).

**Figure 4.**
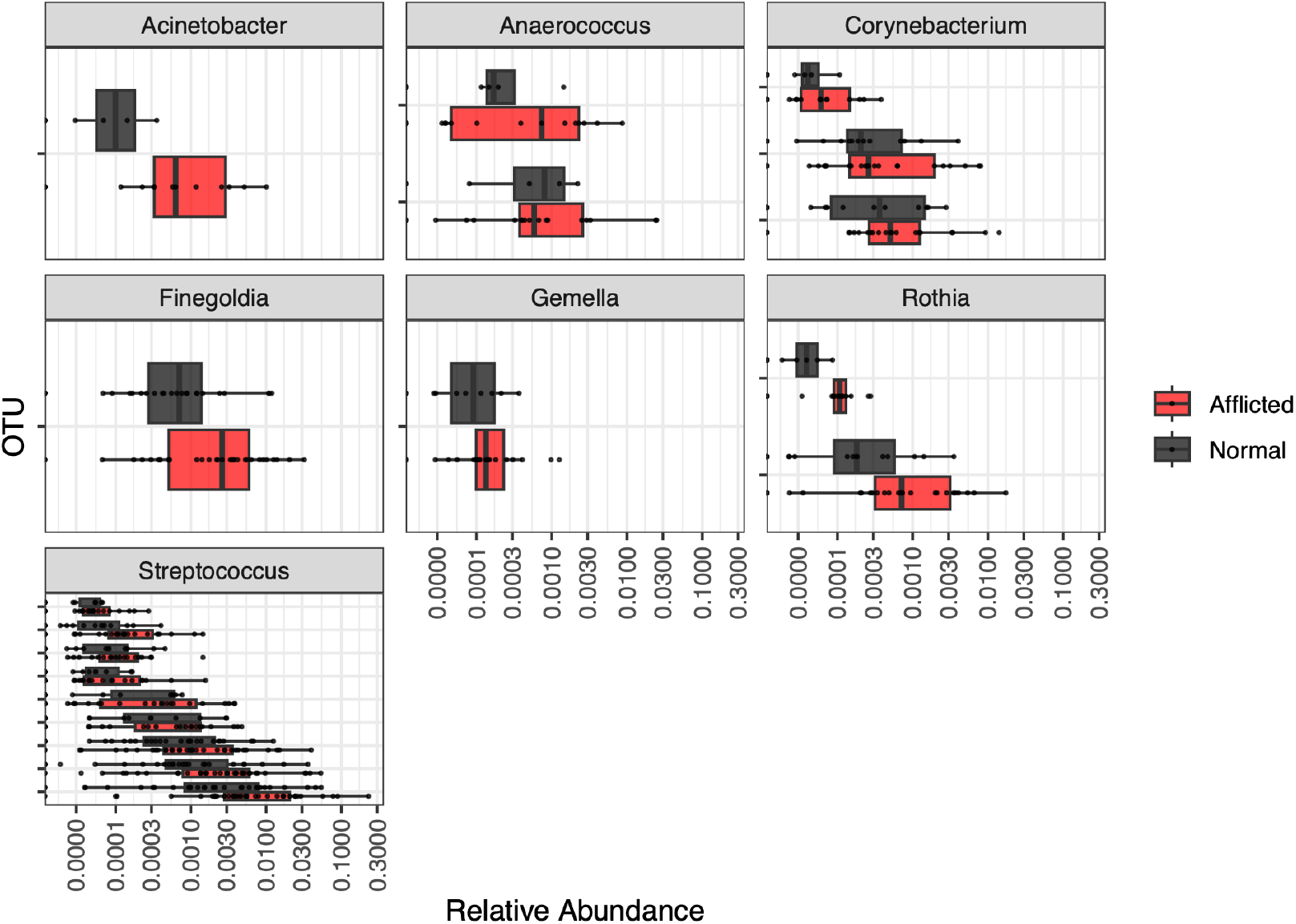
Relative abundances of core taxa differentiating dysbiotic from eubiotic hair sites while controlling for age differences. Within each genus, only OTUs that are significantly associated with dysbiosis are displayed.

## Discussion

In this work, thirty-six individuals from an African-American cohort provided scalp swabs from both afflicted (self-reported hair loss) and normal (no hair loss) sites and the microbiome of each sample was characterized by Illumina sequencing of the 16S rRNA gene.

### Dysbiosis Index

Dysbiosis has been broadly defined as a condition differing from the healthy or normal state [10], however, a consensus on which organisms are harmful has yet to be established. The term dysbiosis is often invoked when describing microbiome changes correlated with disease. However, its use ranges from a broad shift in the microbial community composition to a change in a specific taxon [9]. In addition, dysbiosis is readily applied to the gut microbiome and, therefore, is frequently equated with decreased microbial diversity [9]. This conflation with decreased diversity may lead to confusion, as increased microbial diversity on the skin is a commonly observed with aging and dermatologic disease [15, 32, 33]. In previous work on the scalp microbiome, dysbiosis has been defined as a relative change in ratios of abundant scalp microbiota, such as changes in ratios of *Cutibacterium, Staphylococcus*, and/or *Corynebacterium* species have been reported as markers of dysbiosis [13, 34–36]. More recently, studies have reported that an overall shift in the total microbial community composition, rather than a focus on specific species or taxa, may be a more accurate description of dysbiosis (Table I; [15, 37]).

**Table I.** Summary of findings from studies on the association between microbial dysbiosis and scalp-related dermatological conditions. Note difference between number of health and disease in this study refers to the number of samples from normal and afflicted sites, respectively. Number of samples not identical due sample loss from quality filtering.

| Ref | Populati<br>on | #<br>Healt<br>hy | #<br>Disea<br>se | Condition | Definition of<br>dysbiosis | Diversity | *Staphylococ*<br>*cus* | *Corynebacte*<br>*rium* | *Propionibacte*<br>*rium/*<br>*Cutibacterium* |
| --- | --- | --- | --- | --- | --- | --- | --- | --- | --- |
| [34] | Italian | 15 | 15 | Alopecia<br>Areata (AA) | Increase in ratio of<br>in Cutibacterium<br>(*C. acnes*) to<br>Staphylococcus<br>(*S. epidermidis*, *S.*<br>*aureus*) from<br>qPCR quantified<br>species | Increase | Decrease in<br>Genus<br>Decrease in<br>*S.*<br>*epidermidis* | No change | Increase in<br>Genus<br>Increase in<br>*P. acnes* |
| [35] | Not<br>specified | 12 | 23 | AA<br>cicumscripta,<br>Frontal<br>Fibrosing<br>Alopecia,<br>Lichen<br>Planopilaris | Increase in<br>Firmicutes to<br>Actinobacteria<br>ratio | No<br>Change | Increase | Decrease in<br>Phylum | Decrease in<br>Phylum |
| [13] | Japanes<br>e | 63 | 55 | Androgenetic<br>Alopecia<br>(AGA) | Increase in ratio of<br>Cutibacterium to<br>Corynebacterium | No<br>change | Increase | Decrease | Increase |
| [36] | Korean | 12 | 33 | AA | Increase in<br>Corynebacterium<br>and Cutibacterium<br>relative to<br>Staphylococcus<br>caprae | Increase | Decrease in<br>Genus<br>Decrease in<br>*S. carprae* | Increase | Increase in<br>severe cases |
| [37] | Korean | 46 | 95 | AGA | AGA may be<br>affected by<br>multiple bacteria and caused by dysbiosis of the scalp microbiome, rather than individual bacteria | Increase | Decrease in<br>men only | Increase in<br>men only | No change |
| [15] | Chinese | 51 | 35 | AGA | Machine learning classification using overall combined bacterial and fungal community | No Change | No change | Increase | No Change |
| This study | African American / Black | 32 | 35 | Hair loss | The microbial composition of the sample is closer to the disease group centroid than the normal group centroid as calculated by the dysbiosis index. | Increase | Increase in several OTUs<br>Decrease in *S. epidermidis* | Increase in several OTUs | Decrease in *P. acnes* |

The taxonomic composition of the skin microbiome is a result of a multitude of factors including host physiology, environment, lifestyle, and underlying pathobiology [38, 39]. Therefore, “normal” is subject to interindividual variation. Here, we developed a microbial dysbiosis score as a proxy for scalp health to avoid *a priori* selection of dysbiotic organisms and to account for interindividual and potential cross study and disease model variation. In this approach, we first set calculate the mean microbial composition of each group (afflicted or normal) and then determine whether each individual sample is closer to the afflicted or normal mean composition. This allows for a gradient approach to characterizing dysbiosis so researchers can quantify the degree to which a particular sample differs from normal and set different thresholds depending on study goals.

Using our calculated dysbiosis scores, we determined that 24 of 35 (68.5%) afflicted samples (sparse regions of the scalp) were colonized by microbial communities that were sufficiently different from normal to warrant classification as dysbiotic, while 30 of 32 (93.75%) normal samples were classified as non-dysbiotic. As our cohort was self-controlled (the same participants provided both normal and afflicted samples), this approach is sensitive to intra-individual variation and can be used to assess microbiome changes following cosmetic or medical interventions for hair loss.

### Core taxa contributing to dysbiosis

We further identified which taxa were most important in the development of dysbiosis using a random forest machine learning model and a negative binomial mixed effects model (NBMM) to control for the age variable. Both age and sex account for differences in microbiome composition according to our beta-diversity analyses; however, we did not attempt to control for the sex variable as it is not known to be associated with dysbiosis. Old age, on the other hand, is known to be associated with hyper-diversification and dysbiosis [33, 40]. Our results indicate that an increase in certain OTUs within the genus *Corynebacterium* as well as several less common residents of the scalp, is associated with a dysbiotic scalp microbiome. Taxa within the genera *Acinetobacter, Anaerococcus, Corynebacterium, Finegoldia, Gamella, Haemophilus, Rothia*, and *Streptococcus* were identified as the most significantly associated with dysbiosis (Fig. 4). Several of these genera can be highly pathogenic though the pathogenicity is species dependent. The genera *Acinetobacter* and *Anaerococcus* have previously been identified as more abundant in the scalp dermis of patients with Alopecia Areata [34]. *Streptococcus*, which is implicated in many infectious diseases including cellulitis and toxic shock syndrome [41], was the most significant contributor to dysbiosis in both the RF (Fig. S1).

Both increases and decreases in *Corynebacterium* have been associated with scalp-related conditions (Table I). When we assessed the genus *Corynebacterium* in aggregate, we observed that this genus was more abundant in normal hair sites (Fig 1). However, when the OTUs were assessed individually with a NBMM, some genera were significantly more abundant in the afflicted sites, and some were more abundant in the normal sites (Fig S2), potentially accounting for the discrepancies between previous studies (Table I). In addition, although 446 unique *Corynebacterium* OTUs were identified in the samples, only 3 OTUs contributed to dysbiosis (Fig 4).

Similarly, taxa of the genera *Staphylococcus* were more abundant in normal sites when evaluated in aggregate (Fig 1), however, some OTUs within these genera were more abundant in hair loss sites (Fig 4). Therefore, we recommend that future work evaluate taxonomic abundances without aggregating sequence data at the genus level. A better approach would be to conduct full-length 16S sequencing and/or metagenomic sequencing to identify species-level taxonomy, however, in the case that only hypervariable region 16S data is available, Amplicon Sequence Variation (ASV) or OTU level taxonomic composition would be preferred.

Our findings show that the perception of hair loss is related to differences in the scalp microbiome. These differences may be explained by changes in the scalp microenvironment that result from the loss of hair. Previous work has shown that cutaneous body site characteristics (moisture, sebum content) account for compositional differences in the microbiome composition [42]. Alopecia has been linked to increases in sebum production [43], leading to microenvironment changes that can influence scalp microbiota. Skin-associated bacterial isolates show differing growth profiles in response to both sebum and sweat [44]. Therefore, differences in the scalp microenvironment due loss of hair, including changes in moisture retention, sebum concentration, and even UV exposure may ultimately explain the observed microbiome differences.

Previous work has also shown that hair shafts promote the growth of gram-negative biofilms while inhibiting the growth of gram-positive *Staphylococcus* species [8]. We observed a decrease in *S. epidermidis* OTUs in the afflicted sites, which could have allowed the colonization of other potentially harmful species. In addition, antimicrobial proteins (lysozyme and histones) have been extracted from human hair [45]. Therefore, the loss of hair in the afflicted sites may have opened the door to enhanced colonization and increased diversity due to the loss of protective biofilms and/or antimicrobial properties of the hair, potentially exacerbating microbiome differences. Some degree of protection from colonization of harmful bacteria was seen in younger individuals, as microbial diversity only increased significantly at the afflicted sites for those participants at or above the median age of 52 years (Fig. 1). Alternatively, the older individuals in the cohort may have had differing underlying scalp disorders. Recent work has found microbial dysbiosis was not confined to areas of hair loss, rather it extended across the scalp in male subjects with androgenic alopecia [15]. Therefore, the baseline group in our study may have also exhibited dysbiotic features. Taken together, these results indicate that changes in the scalp microbiome are likely a consequence of both underlying pathobiology and further exacerbated by physical microenvironment changes.

### Limitations

While this study offers a unique method to qualify scalp health, certain limitations present challenges for interpretation. This work does not include formal clinical diagnoses, which may explain why only 68.5% of the afflicted sites were classified as dysbiotic, as there may be a higher degree of variance and underlying mechanisms within the afflicted group of samples. For example, androgenetic alopecia is more common in older individuals, and this condition may elicit a different microbial response compared to traction alopecia, which is the predominant alopecia type in younger individuals of African descent [46]. Hair loss can also be due to a variety of factors such as stress, nutrient deficiencies, or hormonal changes that are unrelated to underlying pathobiology. In addition, participants were instructed to collect their own scalp swabs and participants were not given specific instructions regarding washing prior to the swabs, which introduces another layer of variability and bias to the measurements. Ideally, unbiased measurements of hair density should be reported with imaging-based techniques.

Despite these limitations, our study highlights that self-perceived hair loss results in a measurable and statistically significant difference in the scalp microbiome. Future work should consider underlying pathobiology, inclusion of a control group with no evidence of hair loss, imaging-based assessments of hair-loss, as well as incorporating a larger number of subjects and range of ethnic groups, as most studies to date focus on a single ethic group.

## Conclusions

This study demonstrates that self-perceived hair loss in an African American cohort is associated with significant and measurable alterations in the scalp microbiome. We have established that these subjective consumer assessments correspond directly to objective changes in both microbial community diversity and composition. Furthermore, we have developed a quantitative dysbiosis index that effectively captures the degree of microbial deviation from a normal, healthy scalp state. Our analysis not only confirmed the presence of dysbiosis in affected areas but also identified a core set of operational taxonomic units (OTUs) from genera *Acinetobacter, Anaerococcus, Corynebacterium, Finegoldia, Gamella, Rothia*, and *Streptococcus* as key drivers of this dysbiotic state. Notably, the differential abundance of these taxa was highly specific at the OTU level, underscoring our finding that aggregating data at the genus level can obscure critical biological signals and is not recommended for such analyses. The dysbiosis index presented here addresses a critical need in cosmetic science by providing a quantitative tool to assess product efficacy by measuring a direct biological response. By bridging the exceptional gap between a subjective consumer experience and an objective microbiological metric, this index provides a potential way to evaluate the impact of cosmetic interventions on scalp health. Future research should focus on validating this index across broader populations and exploring treatment modalities designed to specifically modulate these dysbiosis-associated taxa. Ultimately, this work paves the way for a new generation of targeted, microbiome-informed strategies for managing hair loss and improving consumer well-being.

## Data Availability

The data underlying this article are available in Figshare (10.6084/m9.figshare.28829897). Associated code is available at https://github.com/jud-m/Defining-microbial-dysbiosis.

