## Supplemental Figures for "Scalp microbiome differences in subjects with self-reported hair loss: A quantitative approach to microbial dysbiosis"

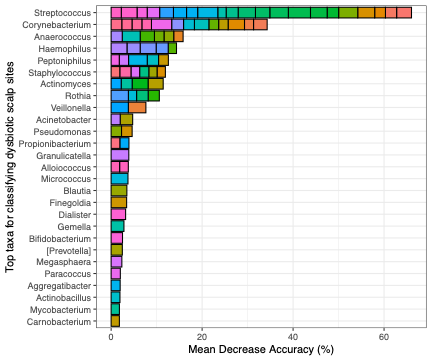
**Supplementary Figure 1**. Mean decrease accuracy of top 100 OTUs for classifying dysbiotic scalp sites as determined by a random forest (RF) classifier. Stacked bars represent

individual OTUs within each genus.

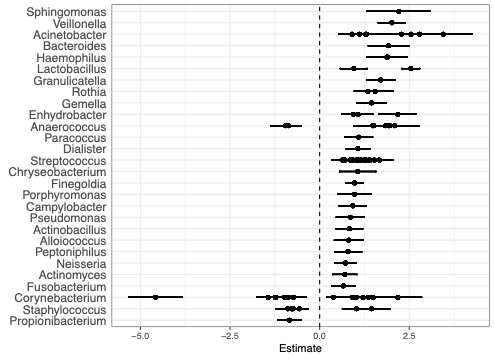

**Supplementary Figure 2**. Fixed effects coefficients for significantly (p <0.05) differentially abundant taxa as determined by a mixed effects model with area condition (afflicted or normal) as the fixed effect and age as the random effect. The estimate (β) gives the multiplicative

change in the mean OTU count for afflicted sites in comparison to normal sites. Each

point with error bars within a genus represents an individual OTU.
